# An optimal variant to gene distance window derived from an empirical definition of cis and trans protein QTLs

**DOI:** 10.1101/2022.03.07.483314

**Authors:** Eric B Fauman, Craig Hyde

## Abstract

**Background:** A genome-wide association study (GWAS) correlates variation in the genotype with variation in the phenotype across a cohort, but the causal gene mediating that impact is often unclear. When the phenotype is protein abundance, a reasonable hypothesis is that the gene encoding that protein is the causal gene. However, as variants impacting protein levels can occur thousands or even millions of base pairs from the gene encoding the protein, it is unclear at what distance this simple hypothesis breaks down.

**Results:** By making the simple assumption that cis-pQTLs should be distance dependent while trans-pQTLs are distance independent, we arrive at a simple and empirical distance cutoff separating cis- and trans-pQTLs. Analyzing a recent large-scale pQTL study (Pietzner, 2021) we arrive at an estimated distance cutoff of 944 kilobasepairs (kbp) (95% confidence interval: 767–1,161) separating the cis and trans regimes.

**Conclusions:** We demonstrate that this simple model can be applied to other molecular GWAS traits. Since much of biology is built on molecular traits like protein, transcript and metabolite abundance, we posit that the mathematical models for cis and trans distance distributions derived here will also apply to more complex phenotypes and traits.

## Background

Genome-wide association studies (GWAS) have been highly successful at identifying reproducible and robust genetic associations for a wide variety of human phenotypes. However, because GWAS identify loci and not genes, the identity of the gene mediating the impact on the phenotype is not always clear.

When the phenotype being considered is the abundance of a particular protein a natural hypothesis is that the gene encoding the protein (the cognate gene) is the causal gene, that is DNA variants that correlate with interindividual differences in protein abundance are somehow influencing the gene encoding the protein resulting in differences in levels of the protein. Such a DNA variant is termed a protein quantitative trait locus, or pQTL.

Many GWAS of protein abundance have now been conducted generating tens of thousands of published pQTLs. The genomic position of a pQTL for a specific protein is often located near the cognate gene. As a practical matter, a pQTL near a cognate gene is called a “cis-pQTL” with the assumption that that pQTL is acting through the cognate gene. When a pQTL is located far from the cognate gene (or on a different chromosome), this is called a “trans-pQTL” with the assumption that that pQTL is acting through an intermediate gene, for example a transcription factor near the trans-pQTL which then drives expression of the cognate gene. Various distance cut-offs have been employed to segregate cis-pQTLs and intrachromomsomal trans-pQTLs, with a distance of 500,000 bp as a typical value.

In order to arrive at an empirical definition of cis- and (intrachromosomal) trans-pQTLs we took a closer look at the distribution of variants to cognate gene from a recent large scale pQTL study [1]. We start with the simple assumption that for cis-pQTLs there should be some (non-random) distance dependence between the variant and the transcription start site (TSS) of the cognate gene, while for trans-pQTLs, there should be a random distribution of distances. From these assumptions, we build a simple model that fits the distribution of intrachromosomal variant-TSS distances across the full range of observed distances.

## Results

Pietzner et al identified 2,051 intrachromosomal pQTLs [1]. The variant-TSS distances for these pQTLs fall into 33 log-based distance bins (width = 10^0.25^). The histogram of these log-transformed distances shows two distinct peaks, the first with 251 pQTLs in the bin covering distances from 13,335 to 23,713 bp, and the second with 84 pQTLs in the bin covering distances from 74,989,421 from 133,352,143 bp (Figure 1).

**Figure 1:**
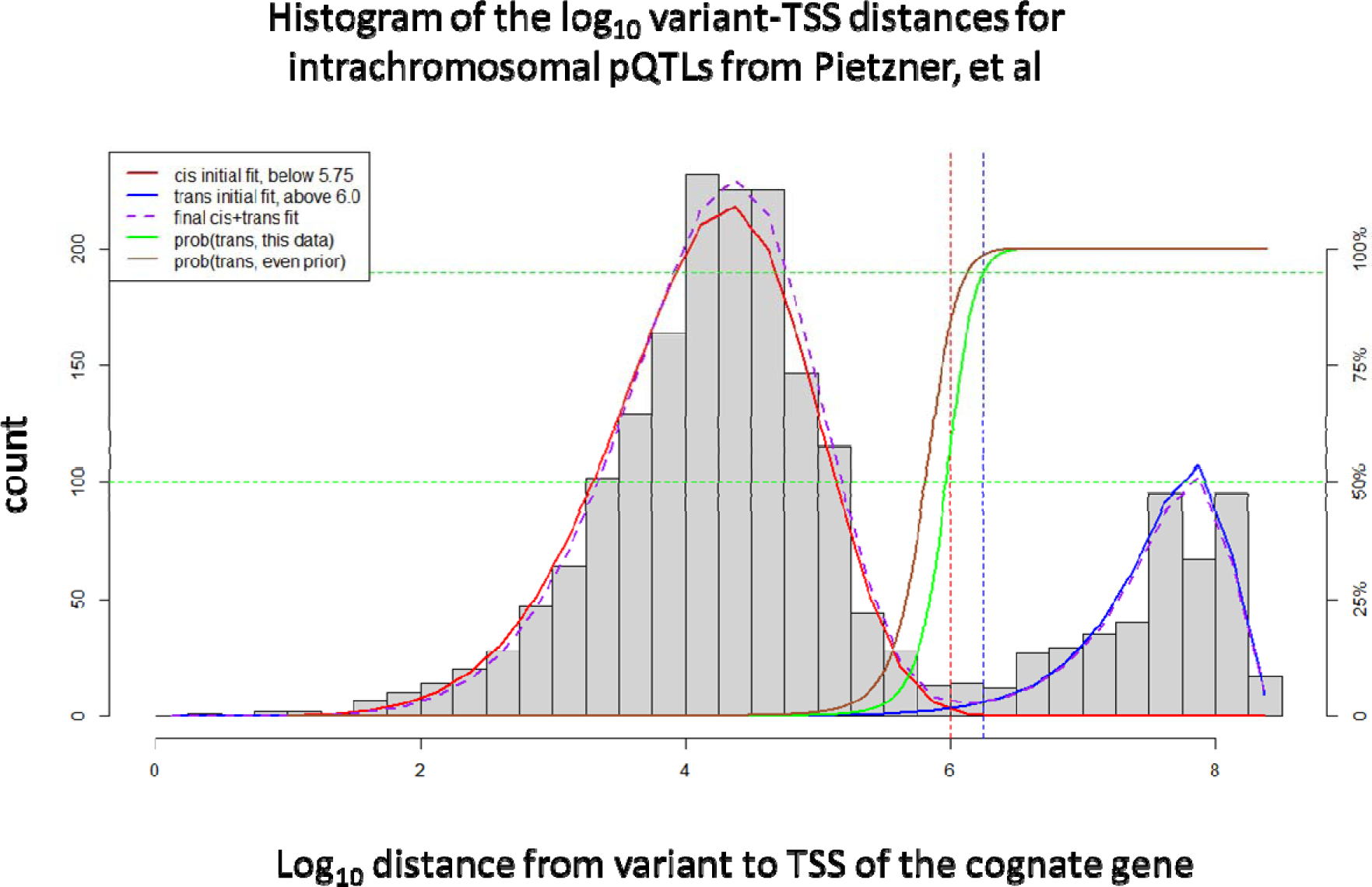
Histogram of log base 10 of the distance from lead SNP for GWAS of protein abundance to the transcription start site (TSS) of the cognate gene for that protein, for 2051 unique proteins from the study of Pietzner, et al. Four bins are used for each log unit. Solid red line represents the best fit Weibull distribution curve fit to all data points below 10^5.75^. Solid blue line represents best fit random distribution curve fit to all pQTLs with a distance beyond above 10^6^ base pairs. Dashed purple line represents best combined model starting from the parameters estimated for the initial Weibull curve and adding a Weibull fraction parameter to add the Weibull curve and the trans model curve.

The second peak (longer distances) follows the expected distribution for two points selected at random from the human genome within the same chromosome, which is to say there is no explicit distance dependence on the variant-TSS distances aside from the physical requirement that the two positions occur on the same chromosome. This distance-independent distribution requires no free parameters. The first peak (shorter distances) is well fit by a two-parameter Weibull distribution, which represents a specific model for the distance dependence on the distribution of variant-TSS distances. The full model requires a third parameter which is the proportion of the observations falling in to the first peak or the second peak. It should be noted that when plotted using the untransformed variant-TSS distance the distribution resembles a rapidly decreasing exponential decay, with a maximum density close to 0 (Figure 2).

**Figure 2:**
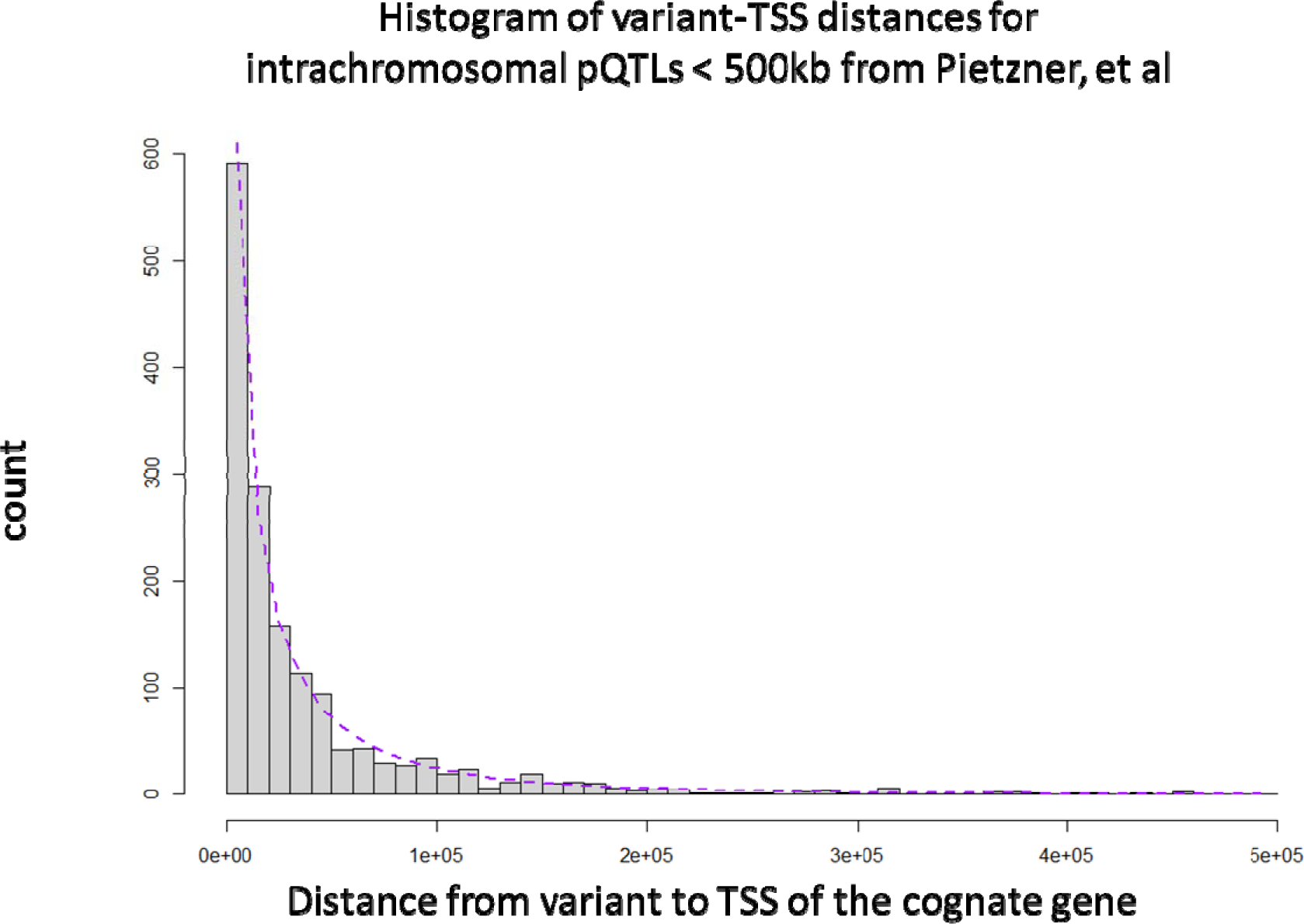
Histogram of distance from lead SNP for GWAS of protein abundance to the transcription start site (TSS) of the cognate gene for that protein, for 1604 unique proteins where the distance is less than 500 kb (bin size = 10 kb), with the curve fit to our global model which includes a Weibull curve and our trans model. The Weibull model dominates at distances less than 1,000,000 base pairs.

The full distribution is well fit by our combined model. The three parameters (and the standard errors) for the full model are *κ* (Weibull shape parameter) = 6.78 (0.211), *λ* (Weibull scale parameter) = 4.48 (0.024), and Weibull fraction = 0.799. The one-sample Kolmogorov-Smirnov test for this fit yields a p-value of p=0.36, consistent with this simple model being a reasonable explanation for the observed distribution. By contrast, using instead a two parameter (mean and sd) Gaussian to fit the left-most peak in a similar three parameter model yielded an analogous Kolmogorov-Smirnov test p-value of p = 0.005, indicating a significantly poorer fit.

Using these parameters from the best-fit Weibull-based model, we calculated where the regimes cross-over, that is, the distance at which the Weibull and the random model are equally likely. For this data set, with ∼80% Weibull fraction, that cross-over occurs at a distance of 944 kilobasepairs (kbp) with a 95% confidence interval of [767 – 1,161] kbp. If we assume a model where the two regimes are *a priori* equally likely, that cross over would occur at 653 kbp, with a 95% confidence interval of [556 – 767] kbp.

A separate large-scale pQTL study using the same platform but applied to a different population was recently published [2]. Ferkingstad et al identified 18,084 associations across 4,907 Somascan aptamers in their study of 36,000 Icelandic subjects. Filtering to the strongest intrachromosomal pQTL per aptamer we derived 2,315 variant-gene distances ranging from 1 bp to 224,221,289 bp. For this data set, the full combined model of yielded values of *κ* = 6.19 (0.315), *λ* = 4.47 (0.043), and Weibull fraction = 0.665. The one-sample Kolmogorov-Smirnov test for this fit yielded a p-value of p=0.022, consistent with this simple model being a reasonable explanation for the observed distribution. Again, the analogous fit using a Gaussian instead of a Weibull for the left-hand peak yielded a poorer fit, with p=0.004 for the Kolmogorov Smirnov test. For this data set, the regime cross-over point occurs at 966 kbp, with a 95% confidence interval of [700 – 1,303] kbp.

Another genetic trait where the cognate gene is a reasonable hypothesis for the causal gene is mRNA abundance, in which case the genetic variant represents an expression quantitative trait locus (eQTL). The eqtlgen [3] study represents a very large, well powered eQTL GWAS. To reduce the computation involved this study only calculated association statistics for variants within 1 Mb of the midpoint of the cognate gene, limiting our ability to use this study to analyze long-range intrachromosomal eQTLs. Despite this truncation, the distances under 1 Mb have a reasonable fit to the Weibull distribution, with shape and scale parameters similar to those observed in the two pQTL studies (of *κ* = 6.19 (0.315), *λ* = 4.47 (0.043))

When the genetic trait is metabolite abundance (metabolite quantitative trait locus, or metabolite QTL) the known biochemistry can point to a likely causal gene [4]. Using all available metabolite QTLs in the GWAS catalog paired with a large set of curated metabolite interacting proteins [5] we identified 250 intrachromosomal variant-gene pairs, of which 53 have a variant-TSS distance of less than 1 Mb. These variant-TSS distance can also be fit with our combined model, with Weibull shape and scale parameters of *κ* = 4.262(1.881) and *λ* = 5.544 (0.708), respectively. For this analysis the Weibull fraction is only 35%, probably indicating that the limited number of curated metabolite interacting proteins is missing many of the true causal genes.

## Discussion

The GWAS catalog now contains over 300,000 genetic associations, but for the majority of these the underlying causal gene, the gene mediating the phenotypic impact of the genetic variation, is unknown. While the genes close to the genetic variation often represent plausible candidate genes, a precise definition of “close” has been difficult to define.

By relying on molecular traits which minimize the assumptions involved in selecting causal genes we have been able to identify two populations of variant-gene distances; one population where the distribution of distances is a function of the distance of the gene from the variant, and a second population where the distances are dictated by the mathematics of picking two points at random. The first population follows a Weibull distribution and is substantially contained within the interval from 0 to 1 Mb. For the second distribution, because individual chromosomes are over 100 Mb long, two randomly selected intrachromosomal points are almost always (99%) more than 1 Mb apart. Thus, these two populations are well-separated and can be interpreted as the mathematical representations of the biological processes of cis- and transQTLs.

Previous analyses of molecular QTLs have similarly noted a rapid drop-off of observed associations with increasing variant-TSS distance. For example, Roby Joehanes et al used a multi-exponential decay with a median variant-TSS distance of 27 kb to model a large set of eQTLs measured in whole blood samples from over 5000 participants in the Framingham Heart Study [6]. There is however no theoretical model to rationalize an exponential decay for this distribution.

As noted by Lieberman-Aiden et al [7], the distribution of promoter-enhancer Hi-C distances can be modeled using a power-law with an exponent of -1, consistent with a Fractal Globule model of a self-avoiding compact polymer [8]. Plotted against variant-TSS distance, a power-law with p(distance) proportional to 1/distance looks similar to an exponential decay, with p(distance) proportional to e^-distance^. However, a simple 1/distance power-law distance dependence would not generate the curve obtained in Figure 1, since power-law would place an equal numbers of observations in each bin, since the bins increase in width with increasing distance at the same rate that p(distance) is decreasing.

The Weibull distribution used here was first described as a family of curves [9] which has found applicability to describe the distribution of particle sizes following fragmentation or fractionation [10]. Brown &Wohletz provide a mechanistic derivation for the Weibull distribution which follows from repeated fragmentation of a larger structure, with each step resulting in a fractal fragmentation pattern (thus following a power law). Smaller fragments escape further fragmentation, resulting in a rapid drop-off of larger particle sizes.

An entirely hypothetical conjecture would be that the pattern we observe in this data results from a similar superposition of multiple processes. The Activity-by-Contact model of enhancer-promoter regulation suggests that the activity of a particular enhancer-promoter pair is increased by the strength (activity) of the enhancer and decreased by the distance between the enhancer and promoter[11, 12]. Since any given promoter can be influenced by multiple enhancers, the strongest genetic associations are more likely to come from closer enhancers. The dense packing of the chromosome provides the equivalent of the single fractionation event, imposing a fractal distance geometry on the genome. The fact that there are far more enhancers than promoters in the genome provides the equivalent of multiple fractionation events, potentially explaining the fit to the Weibull distribution for molecular QTLs in the range of 0 to 1 million base pairs (Figure 4).

Trans-eQTLs and trans-pQTLs are generally understood to be acting on a gene proximal to the variant which then influences the molecular trait of interest. The cis fraction in our combined model is then likely to reflect the extent to which we have correctly selected the set of true causal genes for a given study. Further, the model suggests that in general about 99.9% of GWAS variants should be explainable through a gene with a TSS within 1 megabase of the lead variant. Thus, if a large fraction of the variant-TSS distances fall into the long-range, distance-independent regime our model suggests it is worth taking another look at the set of proposed or potential causal genes.

Assuming that most GWAS variants are likely impacting biology through their influence on molecular traits such as transcript, protein or metabolite abundance we expect that the cis- and trans-models and distributions observed here will apply to other, more complex or polygenic traits. It should be noted however that the exact mechanism linking the GWAS variant to the causal gene is not addressed in this model. It has been observed that a large fraction of pQTLs and metabolite QTLs are linked to missense variants, and that may skew the exact distributions somewhat when looking at other phenotypes or disease traits.

An additional caveat is that this study focused on only the single strongest association per molecular trait (per chromosome) and this will tend to bias the set of variants as well. This simplification was applied here because while it is straight-forward to define the primary signal per locus there are still multiple approaches to defining secondary or independent signals. As molecular QTL studies continue to grow in size and power, it will be important to revisit this analysis with respect to secondary signals.

## Conclusion

By leveraging recent large-scale molecular QTL genetic studies we demonstrate that variant-TSS distances fall into one of two regimes, a short-range, distance-dependent one, or a long-range, distance-independent one. These correspond to the biological notions of cis and trans genetic effects. By providing mathematical models for these two regions we demonstrate a clear separation occurring at about 944 kbp in the situation where 80% of observations are well explained by cis effects, or 650 kbp when cis and trans effects are equally likely.

## Methods

Pietzner et al reported 10,674 unique pQTLs across 3,892 distinct proteins. For this analysis we only used the primary association reported for each locus. Further we eliminated all SNP-protein pairs where the SNP and the cognate gene occur on the different chromosome in the latest build (Ensembl release 104). Variants were converted from the GRCh37 coordinates provided in the original data table by reference to an rsid if provided and current, or via the NCBI Genome remapping service. Proteins were mapped to HGNC gene symbols using the Ensembl gene IDs if provided and current. Otherwise HGNC gene symbols were assigned manually from the provided protein names. As some proteins actually map to multiple genes (either due to complexes or ambiguity), SNP-protein pairs were retained if exactly one of the cognate genes was present on the same chromosome as the SNP. This resulted in a total of 2,051 intrachromosomal SNP-gene pairs, where the distance from the SNP to the TSS of the gene was between 3 and 206,513,449 base pairs.

Given the eight orders of magnitude range for variant-TSS distances we log-transformed the distances (specifically using 10 as the base and binning at 0.25 log_10_ units to generate 4 bins per order of magnitude). For visualization and curve-fitting, we used the number of intra-chromosomal pQTLs falling in each bin based on this log-transformed distance.

Our trans model is a mathematical model of the distribution of two random positions in the genome that happen to fall on the same chromosome. To generate this trans model, we represented all pairs of two randomly selected positions within a single chromosome as an NxN matrix, where N is the length of the chromosome, and rows and columns representing the 1^st^ and 2^nd^ position, respectively. With the exception of a discontinuity at distance = 0 (also not handled by log-transformation, but highly unlikely and not present in our data), the number of random pairs at a distance d > 0 bp is given by the two diagonal segments shifted ‘d’ units from the central diagonal, and so totals 2*(N-d) out of N^2^ elements (where d < N). The probability of distance ‘d’ within the chromosome is then prob=2*(N-d)/ N^2^ if d < N, otherwise prob=0. For any one chromosome then this probability is linear with respect to distance, with a maximum at a distance of 1 and a minimum at a distance corresponding to the length of the chromosome.

Restricting to random pairs on the same chromosome, the relative likelihood of a pair being in any chromosome ‘i’ is given by its relative number of elements to that of all intra-chromosomal pairs on the genome, so that prob(Chr=i) = N_i_ ^2^/ Σ N _i_^2^, where N_i_ is the length of chromosome ‘i’. The final probability of a random intrachromosomal pair having distance d > 0 bp is then given by the weighted sum of the probability at each chromosome, with weights given by the relative likelihood of each chromosome. Hence:

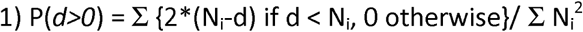

where each sum is over 23 chromosomes, N_i_ being the length of each chromosome. Applied to the observed pQTL data our trans model is only a reasonable fit in the range from 1 to 230 megabase pairs (figure 3).

**Figure 3:**
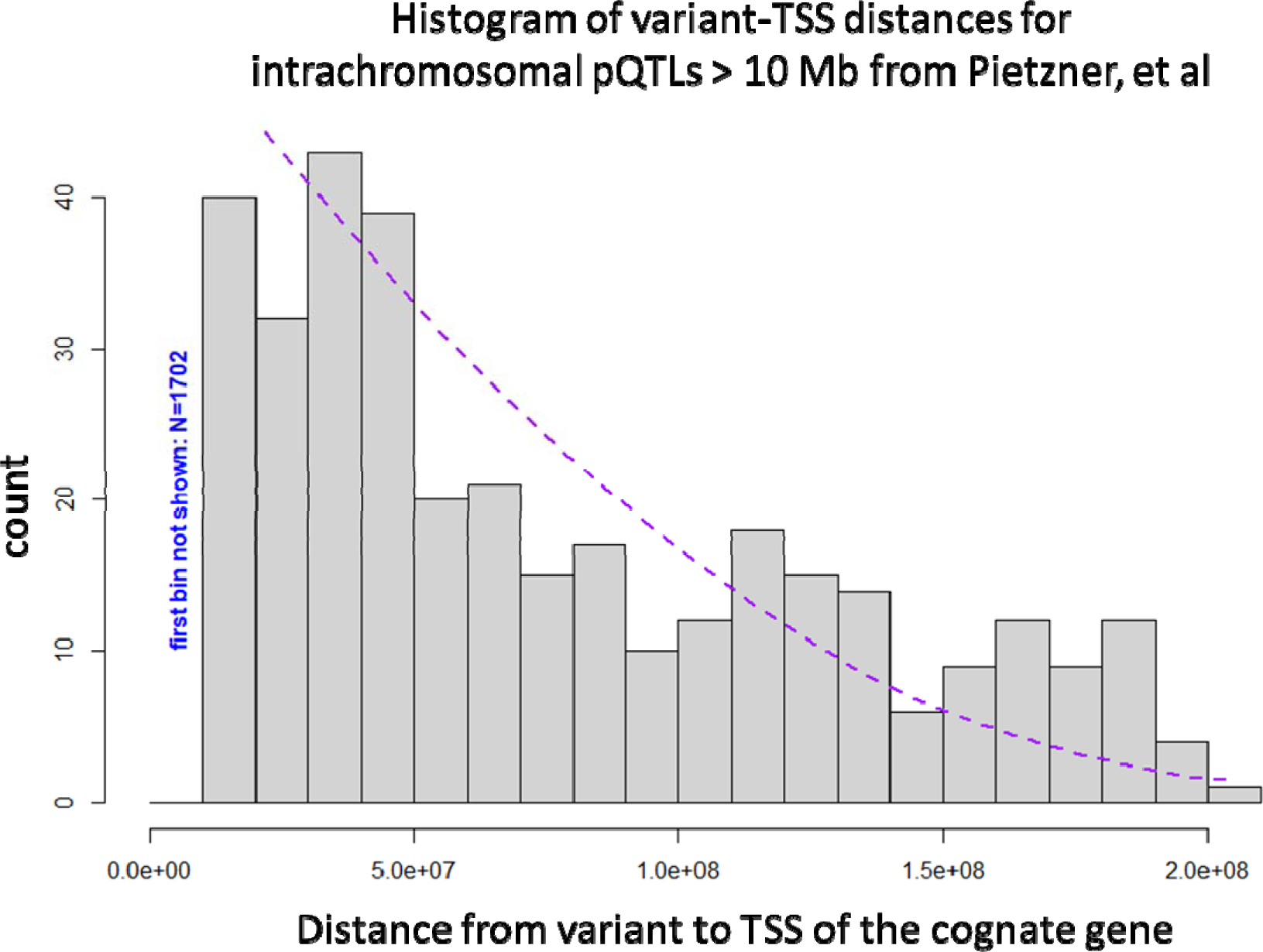
Histogram of distance from lead SNP for GWAS of protein abundance to the transcription start site (TSS) of the cognate gene for that protein, for 349 unique proteins where the distance is greater than 10 megabases (bin size = 10 megabases), with the curve fit to our global model which includes a Weibull curve and our trans model. The trans model dominates at distances past 1 megabase.

**Figure 4:**
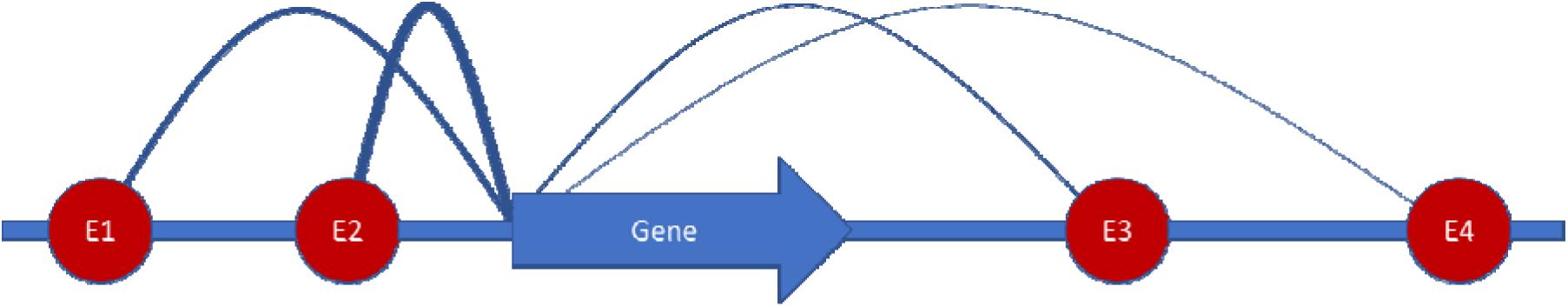
A post-hoc rationale for the Weibull distribution. According to the ABC model [11] of gene activation and the Fractal Globule [8] model of chromatin compaction, the chance that a particular enhancer (E1-E4) is in contact with the promoter of a particular gene (“Gene”) is proportional to 1/distance (that is, distance to the power -1) from the enhancer to the promoter. In a scenario where all enhancers are equally active, a particular gene will be most strongly influenced by the closest enhancer (E2 in this figure). A Weibull model, as observed empirically in this analysis, can result from such a “superposition” of power-law distributions [10].

We found empirically that the remaining (shorter) distances could be fit with the two parameter Weibull distribution. The Weibull distance distribution can be represented as:

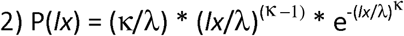

where ***lx*** is the log of distance from the pQTL to the TSS of the cognate gene, *κ* is the shape function and *λ* is the scale parameter.

Combining the Weibull and the random distance models requires one additional parameter defining the relative proportion of pQTLs in a study which fall under into the two regimes.

After establishing that this combined model is a good fit for the Pieztner et al pQTL study we applied the same model to second pQTL study, a large eQTL study and a metabolite QTL study.

For a pQTL study by Ferkingstad et al [2] we used the variant-TSS distance reported in that study’s Supplementary Table 2, column W (distTSS) filtered to the primary hit per chromosome, column AD (Rank) and filtered to the strongest intrachromosomal pQTL per protein, column AC (unadjusted - log10(P), where the TSS is for the cognate gene.

For the eQTL study reported by Vosa et al [3] we used the eQTL with the largest absolute value Zscore (column labeled “Zscore”) for each gene and calculated the variant-TSS distance as the absolute value of the difference between the reported variant position (“SNPPos”) to the TSS associated with the Ensembl gene ID in GRCh37 according to Ensembl for variants within 1 Mb of the TSS of the cognate gene. We only retained associations where the absolute value of the Zscore was at least 5.4485, corresponding to a p-value of less than 5×10^−8^.

To apply the method to metabolite QTLs we first assigned an HMDB identifier [5] to all relevant endpoints reported in the GWAS catalog [13]. We then extracted all “interacting genes” defined for each metabolite as reported in the HMDB. For any metabolite with a single interacting gene on a particular chromosome we identified the strongest reported association for that metabolite on that chromosome. If the reported p-value was less than 5×10^−8^ we calculated the distance from that strongest metabolite QTL to the unique interacting gene.

## Supporting information

pqtl distances and meta data for Ferkingstad

eqtl distances and meta data for eqtlgen

mqtl distances and meta data this work

pqtl distances and meta data Pietzner

---

**Table 1:**
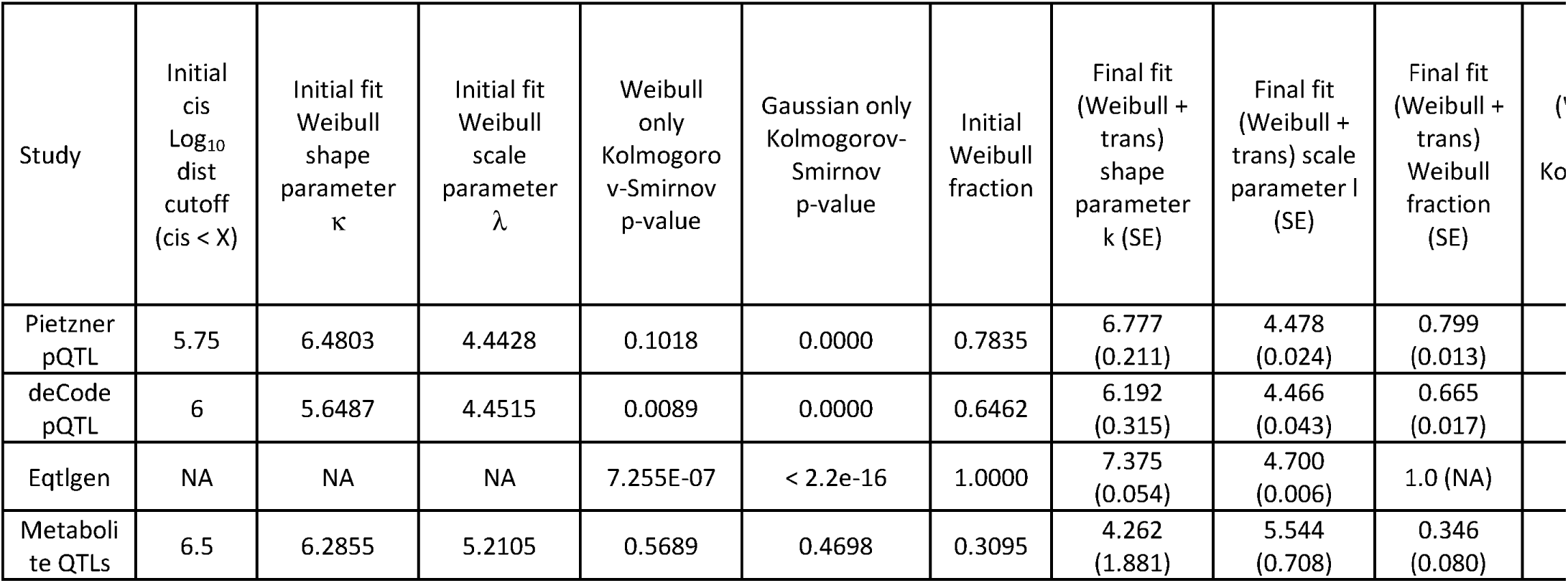
Curve fit parameters for histograms of log_10_(variant-TSS) distances from several studies. Initial fits were from log10=0 to observed minimum in the histogram, as presented in the table. Final fit included our trans model (methods) and started from the values generated from the initial fits. In all cases, the Kolmogorov-Smirnov test favored the Weibull distribution over the Gaussian distribution.

